# Synaptic m^6^A Epitranscriptome Reveals Functional Partitioning of Localized Transcripts for Dynamic Tripartite Synapse Modulation

**DOI:** 10.1101/221374

**Authors:** Daria Merkurjev, Wan-Ting Hong, Kei lida, Belinda J Goldie, Hitoshi Yamaguti, Ikumi Oomoto, Takayuki Ohara, Shin-ya Kawaguchi, Tomoo Hirano, Kelsey C Martin, Matteo Pellegrini, Dan Ohtan Wang

**Affiliations:** Statistics Department, University of California at Los Angeles, 8125 Math Sciences Bldg. Box 951554. Los Angeles, CA 90095-1554; Institute for Integrated Cell-Material Sciences (iCeMS), Kyoto University, Yoshida-Honmachi, Sakyo-ku, Kyoto, 606-8501, Japan; Medical Research Support Center, Kyoto University Graduate School of Medicine, Kyoto 606-8507, Japan; Society for the Promotion of Science (JSPS), 5-3-1 Kojimachi, Chiyoda-ku, Tokyo 102-0083, JAPAN; Undergraduate School of Informatics and Mathematical Science, Kyoto University, Yoshida-Honmachi, Sakyo-ku, Kyoto 606-8501 JAPAN; Graduate School of Biostudies, Kyoto University, Yoshida-Honmachi, Sakyo-ku, Kyoto, 6068501, Japan; Society-Academia Collaboration for Innovation, Kyoto University, Yoshida-Honmachi, Sakuo-ku, Kyoto, 6068501, Japan; Graduate School of Science, Kyoto University, Kitashirakawa, Sakyo-ku, Kyoto, 6068502, Japan; Department of Psychiatry and Biobehavioral Sciences, Semel Institute for Neuroscience and Human Behavior, University of California at Los Angeles, Los Angeles, CA90095, USA; Department of Biological Chemistry, University of California at Los Angeles, Los Angeles, CA90095, USA; Department of Molecular, Cell and Developmental Biology, University of California at Los Angeles, CA90095, USA; The Keihanshin Consortium for Fostering the Next Generation of Global Leaders in Research (K-CONNEX), Yoshida-Honmachi, Sakyo-ku, Kyotoshi, Kyoto, 606-8501, Japan

## Abstract

A localized transcriptome at the synapse facilitates synapse-, stimulus-, and transcript-specific synthesis of the local proteome in response to neuronal activity. While enzyme-mediated mRNA modifications have been shown to regulate cellular mRNA turnover and translation, the role of these modifications in regulating synaptic RNA has not been studied. We established low-input m^6^A-seq of synaptosomal RNA to determine the chemically modified local transcriptome in healthy adult mouse forebrain and identified 4,329 selectively enriched m^6^A RNA peaks in 2,987 genes, which we refer to as the synaptic m^6^A epitranscriptome (SME). SME is functionally enriched in synthesis and modulation of tripartite synapses, and in pathways implicated in neurodevelopmental and neuropsychiatric diseases. Interrupting m^6^A-mediated regulation via knockdown of reader YTHDF1 in hippocampal neurons alters expression of SME member *Apc*, and causes synaptic malfunctions manifesting immature spine morphology and dampened excitatory synaptic transmission concomitant with decreased PSD-95 clustering and GluA1 surface expression. Our findings indicate that chemical modifications of synaptic mRNAs critically contribute to synaptic function.

## Introduction

Synapses connect billions of neurons into functional neuronal and neuroglial circuits that underlie information processing and drive behavior. Remarkably, synaptic connections are highly dynamic, undergoing continuous synaptogenesis and synaptic elimination throughout development and adulthood,^1,2^. Synaptic plasticity, the ability of synapses to change at the level of their molecular components, structure, and transmission efficiency in response to neuronal activity, is a fundamental feature of neural network architecture and brain function^3^.

At the center of activity-induced synaptic alteration, local protein synthesis plays a key role in consolidating neuronal activity into persistent structural and functional change^4–13^. Aberrant translation at synapses has been linked to autism, fragile X syndrome and other intellectual disabilities, identifying dysregulated local translation as a core pathophysiological mechanism in neurodevelopmental and neuropsychiatric disease^14–17^. In addition to ribosomes, thousands of transcripts are localized to synaptic compartments encoding proteins involved in diverse functions, including synapse- and neuron-specific molecules, as well as components of the protein synthesis and degradation machinery^18^.

Recent advances in understanding specific cell-type contributions to synapse refinement has revealed that glial cells play active roles in signaling and facilitating synapse formation, function, plasticity, and elimination, giving rise to the concept of a “tripartite” synapse that comprises presynaptic and postsynaptic terminals as well as glial cells^19–21^. The mechanisms of mRNA trafficking and local translation extend to astrocytes, where selectively enriched transcripts and translational machinery can be detected in perisynaptic processes^22^. These new findings implicate multicellular contributions to input-specific, synapse-restricted protein synthesis.

Given the significant contribution of local translation to synaptic function both under physiological and disease conditions, an area of active investigation concerns the post-transcriptional regulatory mechanisms that support the rapid, spatio-temporally coordinated alteration of local proteome composition underlying synaptic plasticity. Current models suggest that signatures carried by individual RNAs, such as sequence motifs, secondary and tertiary structures affect specific interactions with RNA binding proteins and microRNA-mediated regulatory networks^23–28^. Recent studies of the cellular transcriptome have focused attention on epitranscriptional regulation via enzymatic modification as a mechanism that can dynamically regulate the fate of individual RNA transcripts^29,30^.

Between 0.1 and 0.4% of all adenosines in mammals are modified by the addition of a methyl group to the N6 position of the adenine to form N^6^-methyladenosine (m^6^A), the most abundant mRNA internal modification. m^6^A is reversibly regulated by methyltransferases (“writers”) and demethylases (“erasers”), and has been shown to affect the splicing, trafficking, stability and translation of individual RNAs^31^. In contrast to A-to-I editing, which alters the coding sequence of AMPA or 5HT receptors to regulate signaling, m^6^A alters the local structure and protein binding properties of the modified RNA^32^. The brain contains the highest level of m^6^A expression among the major organs^33^, and many neuronal genes, including genes involved in locomotion, circadian clock regulation, dopaminergic midbrain circuitry, and consolidation of cued fear memory, contain specific modification sites^34–38^. Although evidence for m^6^A as a key post-transcriptional regulator continues to grow, few studies have been performed in neurons and there has been, to our knowledge, no investigation of its presence in the synaptic compartment. We therefore sought to address whether m^6^A could localize to synapses and regulate the local transcriptome.

Here we report an m^6^A epitranscriptome, which we term “SME”, in the synaptic compartment of the adult mouse forebrain. We report that m^6^A is present in the synaptic transcriptome, selectively modifying diverse transcripts in all cellular domains of tripartite synapses. Although elevated methylation has a negative impact on synaptic distribution and abundance of the methylated mRNA in a general way, select mRNAs are hypermethylated at synapses. Localized hypermethylated mRNAs showed strong functional enrichment in synaptic pathways, in sharp contrast to hypomethylated transcripts. We report nuclear and somatodendritic distribution of m^6^A readers, writers and erasers in cortical pyramidal neurons *in vivo* as well as in cultured primary hippocampal neurons. To interrupt m^6^A-mediated regulation in cultured hippocampal neurons, we knocked down dendritically localized reader YTHDF1. YTHDF1-deficient neurons had aberrant spine morphology, with decreased spine volume and dampened spontaneous excitatory synaptic transmission concomitant with decreased PSD-95 clustering and surface expression of AMPAR subunit GluA1. Translation of at least one dendritically localized, highly methylated SME member mRNA encoding APC, a microtubule plus-end tracking protein, was inhibited by YTHDF1 deficiency, further demonstrating m^6^A-mediated regulation of synaptic gene expression and function.

## Results

### m^6^A RNA modification in adult mouse forebrain synaptosomes

We initially crossed two m^6^A-databases with 12 lists of mRNAs localized to axons, dendrites, and neuropils. This comparison revealed substantial overlap (14.0%~56.7%, Supplementary Table S1), strongly suggesting synaptic m^6^A localization. Next, we isolated a synaptosome (SYN) fraction from the forebrains of adult ICR mice (Supplementary Fig. S1a). SYN was enriched for synaptic proteins (Synaptophysin and PSD-95) and dendritically targeted mRNA of *Camk2a*, and depleted of soma-restricted markers (Supplementary Fig. S1b and S1c). An integrity check confirmed intact rRNA bands in all samples and enriched mitochondrial rRNA bands in SYN fractions (Supplementary Fig. S1d); m^6^A dot-blot detected strong m^6^A immunoreactivity in poly(A) RNA isolated from homogenized tissue (HOM) and pooled F3 and F4 (SYN), but not in an *in vitro* transcribed unmodified RNA (Fig. 1a, Supplementary Fig. S1e). Densitometry measured that m^6^A abundance in SYN was ~50% that of HOM (Fig. 1b, Supplementary Fig. S1e).

**Figure 1.**
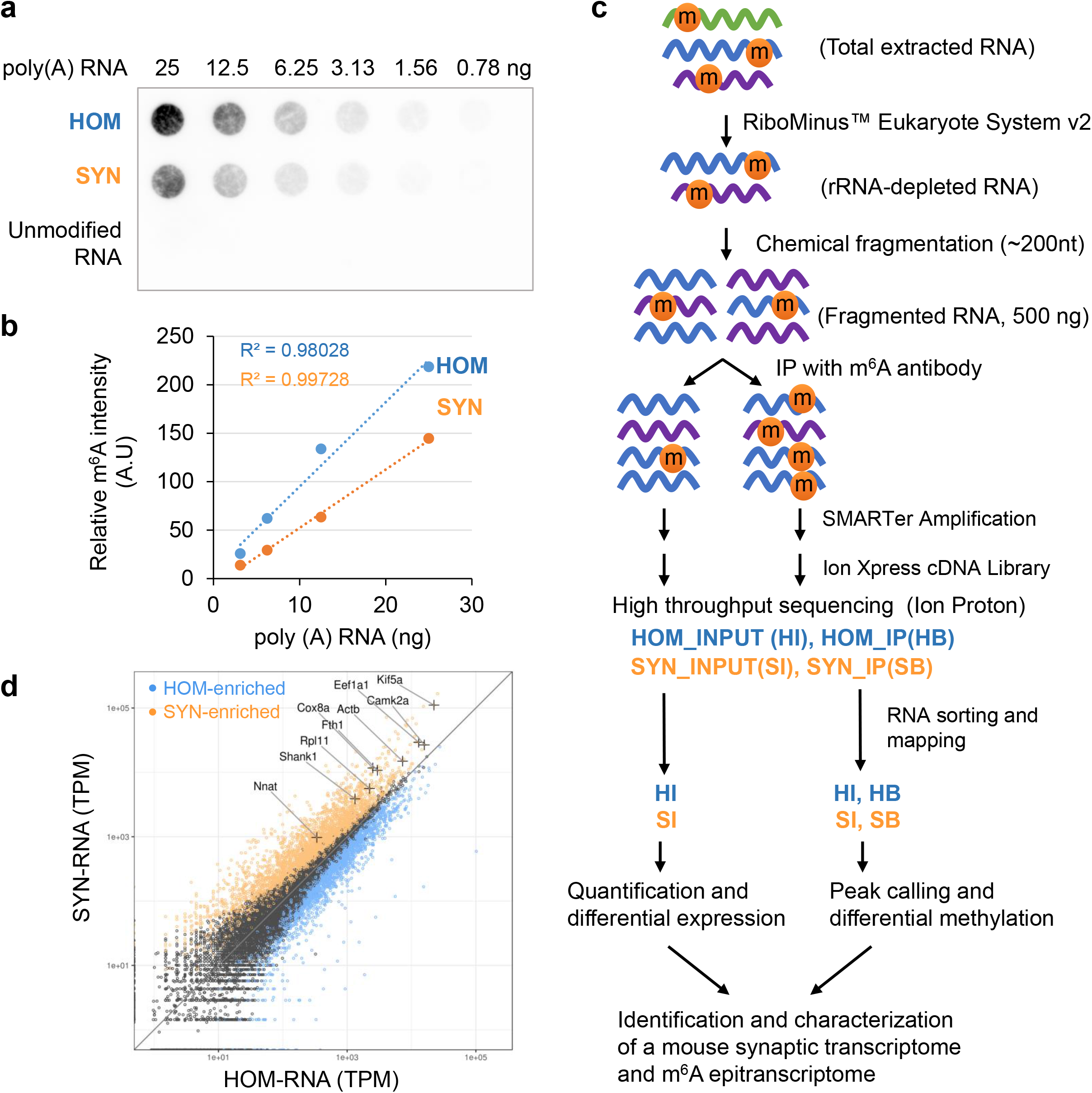
Low-input m^6^A-seq protocol and synaptic enrichment of localized mRNA. a. m^6^A dot-blot of poly(A) RNA purified from HOM (homogenate lysate) and SYN (synaptosomes) starting from 25ng in a twofold dilution series. An equal quantity of *in vitro* transcribed RNA using unmodified nucleotides was used as a negative control. b. Representative dot density plots for the first four dots show linear relationship to the quantity of RNA applied. Approximately 50% m^6^A immunoreactivity was observed in SYN compared to HOM. A.U.: Arbitrary Units. c. Workflow of low-input m^6^A-seq protocol. Four sequencing libraries HI, SI, HB, SB representing HOM input, SYN input, HOM IP, and SYN IP were constructed and characterized. d. Scatter plot comparing HOM and SYN gene expression demonstrates SYN-enriched (orange), SYN-depleted (blue), and genes that are equally represented between HOM and SYN (black). Known localized mRNA species (+) are found in the SYN-enriched gene group, supporting successful enrichment of localized transcripts. TPM: Transcripts per million.

### Low-input m^6^A-seq protocol and synaptic enrichment of select RNA

We then sought to identify and characterize the full synaptic m^6^A-epitranscriptome using m^6^A-seq^39^. The limited yield of RNA from synaptosomes presented challenges using existing m^6^A-seq protocols. Total yield from a single forebrain was approximately 2μg, 1/125th that required by standard protocols^39^. We established a protocol as described in Online Methods to use submicrogram input, by enhancing sample quantification and optimizing IP, and conducting SMARTer amplification prior to cDNA library construction. Our protocol reliably produced INPUT and IP libraries using ribo(-) RNA from 8 mouse forebrains per replicate (Fig. 1c).

Four libraries (INPUT and IP for HOM and SYN) with biological replicates were sequenced (Supplementary Table S2; reproducibility was evaluated using Pearson’s *r* squared, Supplementary Fig. S2). To focus on cytoplasmic RNA species, we computationally removed abundant nucleus-restricted and mitochondrial genome-derived RNA using SortMeRNA (Supplementary Table S3). The remaining uniquely-mapped sequences were used for downstream analysis (Supplementary Table S4). Comparing INPUT libraries (HOM: 17,470 RNA; SYN: 15,767 RNA) revealed 3,067 significantly enriched genes at the synapse (p<0.05 FDR, log_2_FC>0.51) including many known localized mRNA, supporting successful enrichment of synaptic transcripts (Fig. 1d, Supplementary Table S5).

### Identification and characterization of synaptic m6A peaks

We called exon-based m^6^A peaks in HOM and SYN, using recurrent peaks in biological replicates as high-confidence m^6^A sites. We identified 17,009 peaks annotated to 9,600 genes in HOM, and 16,539 peaks annotated to 9,323 genes in SYN (Fig. 2a, Supplementary Table S6). HOM peaks showed high and significant concurrence with previously published peaks in mouse brain tissue^33,40,41^ (Fig. 2b, statistical analysis in Supplementary Fig. S3a), indicating robust and reliable peak detection using the low-input protocol. HOM and SYN peaks showed extensive overlap (Fig. 2c), suggesting a common methylation site selection pathway used by both transcriptomes. The established “RRm^6^ACH” consensus motif (R: A or G; H: A or C or U) was detected in both HOM and SYN m^6^A peaks (Fig. 2d); the distance from peak summit to the discovered motifs was typically less than 100 bases, associating the motif with the antibody binding site (Fig. 2e for SYN peaks, Supplementary Fig. S3b for HOM peaks). In agreement with previous studies, the majority of peaks were detected in CDS and 3’UTR (Fig. 2f). The peak distribution along mRNA showed characteristic accumulation at the STOP codon (Fig. 2g, ^33,42^). Peaks associated with START and STOP codons were identified (Fig. 2h, Supplementary Table S7). Notably, genes with peaks associated with STOP codon, but not START codon, were functionally enriched in human phenotypes of “intellectual disability”, “microcephaly”, “muscular hypotonia”, “hypoplasia of the corpus callosum”, and “seizures,” highlighting their relevance to human brain development and cognitive functions (Fig. 2i and Supplementary Fig. S4; GO analyses using expressed genes in Supplementary Tables S8 and S9).

**Figure 2.**
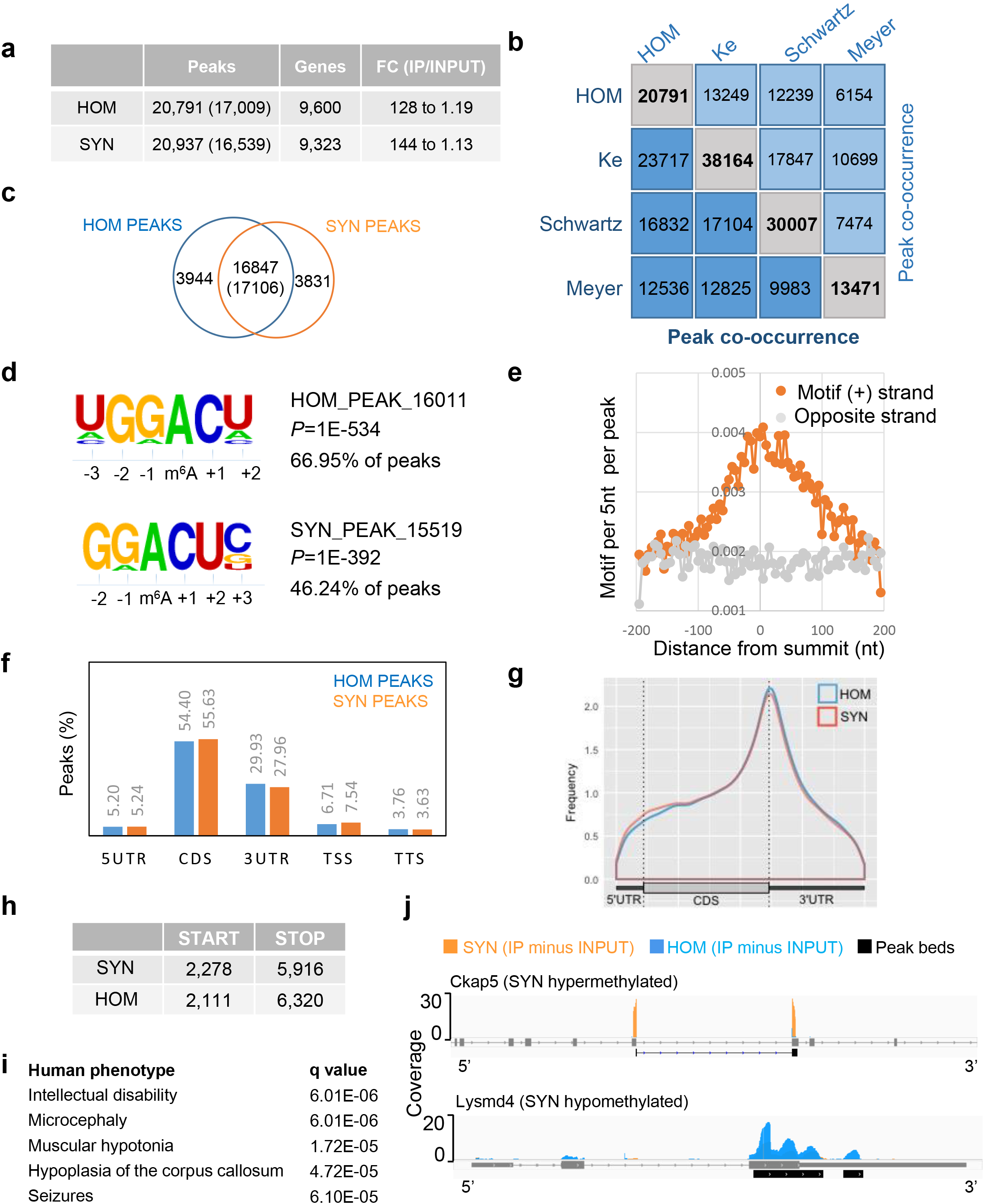
Characterization of m^6^A peaks in HOM and SYN. a. Summary of peaks identified in this study. Numbers in () are counts after joining peaks spanning introns. Fold change represents differential coverage density range between IP and INPUT samples. b. Pairwise comparison of m^6^A peaks identified in HOM to three published studies using mouse brain RNA. The number of peaks identified in each study (grey box), number of overlapping peaks in groups to the left (darker blue), and number of overlapping peaks in groups on the top (light blue). c. Venn diagram of peaks identified in HOM and SYN reveals extensive overlap. () indicates co-occurrence in SYN PEAKS. d. *De novo* motif discovery identified m^6^A consensus motif “RRACU” in both HOM and SYN peaks with variation in adjacent nucleotides. e. Frequency plot of motif per 5nt per peak (y axis) against distance from peak summit (x axis) in SYN. f. Percentage of peaks in HOM and SYN associated with annotated transcript features. TTS: transcription termination site; UTR: untranslated region; TSS: transcription start site. g. Metagene profiles depicting relative frequency of peak coverage along transcripts in HOM and SYN. h. Number of peaks associated with START and STOP codons in HOM and SYN. i. Enriched human phenotypes among genes with stop codon-associated SYN peaks (ToppGene). j. IGV view of peaks in representative synaptically hypermethylated and hypomethylated genes. Ckap5: cytoskeleton associated protein 5; Lysmd4: LysM domain containing 4.

We identified peaks specifically associated with SYN or HOM. Visualization with IGV, shown in Fig. 2j and Supplementary Fig. S5, consistently confirmed differential methylation of RNAs at synapses.

### Methylation negatively impacts synaptic location and abundance of transcripts

We next investigated the relationship between methylation and synaptic RNA concentration, observing weak but significant negative correlation between transcript abundance and methylation (Supplementary Fig. S6a). Furthermore, the concentration of transcripts at synapses relative to whole cells was also negatively correlated with methylation (Supplementary Fig. S6b). This result is consistent with the overall decreased m^6^A level in synaptosomal mRNA (Fig. 1a), suggesting there may be a combinatorial result of local degradation, and retention of m^6^A-mRNA in the soma. Interestingly, methylation in the 5’UTR was not significantly correlated with the synaptic location of transcripts (slope −0.02, Pearson’s *r* −0.02, p value 0.073). This position-specific feature may explain the elevated peak frequency in 5’ region of synaptosomal mRNAs (Fig. 2g).

### A synaptic m^6^A epitranscriptome (SME)

Defining that synaptic m^6^A RNA should not only be detected in SYN, but also enriched in SYN, we next aimed to identify the full synaptic m^6^A epitranscriptome. We merged all m^6^A peaks in HOM and SYN and scored the coverage in the merged peak regions. After normalization to transcript length, 4,329 significantly enriched peaks in SYN in 2,987 genes were identified (p<0.05 FDR, log2FC>0.14, Fig 3a, Supplementary Table S10). We refer to these methylated transcripts as the synaptic m^6^A epitranscriptome (SME).

**Figure 3.**
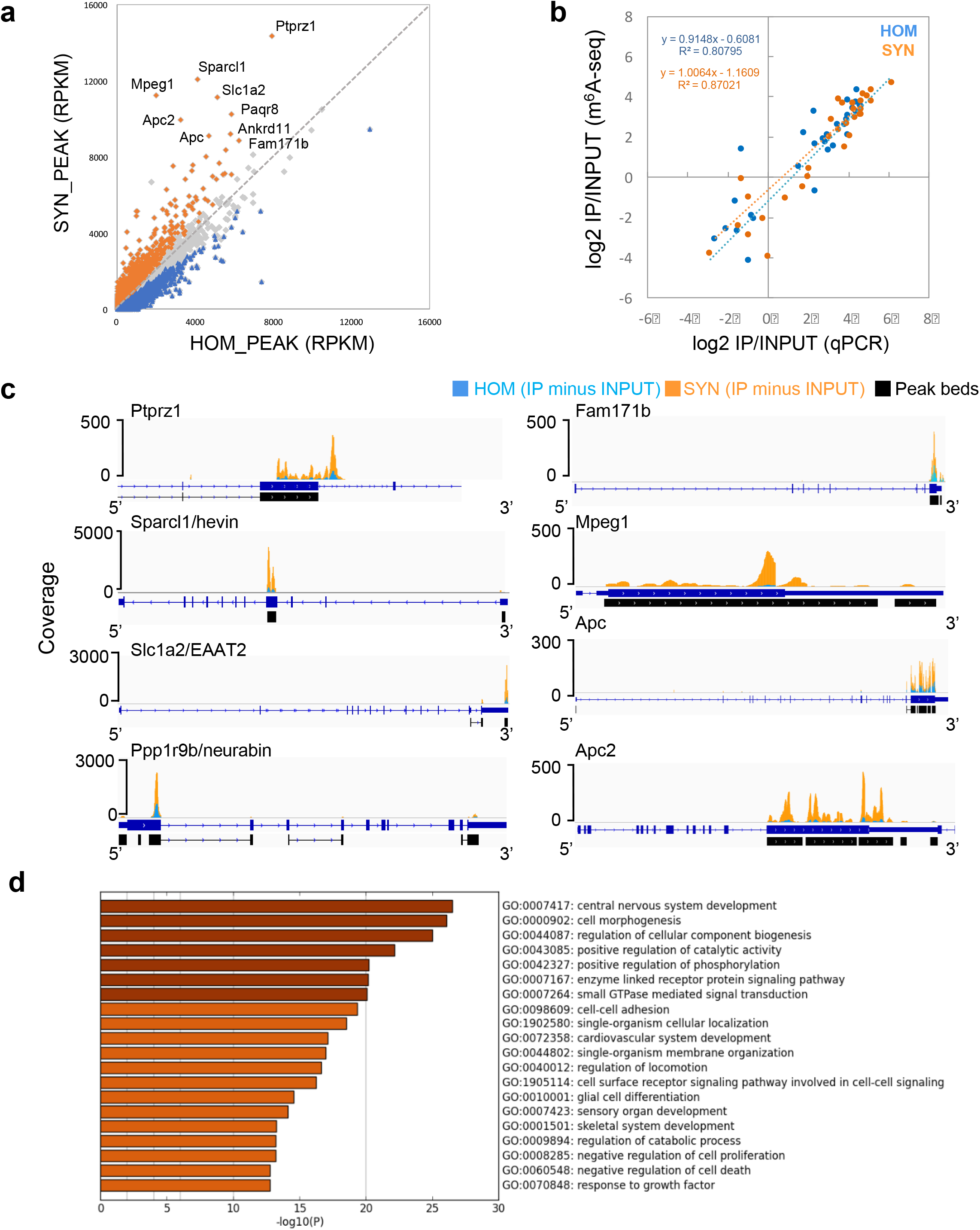
A comprehensive synaptic m^6^A epitranscriptome. a. Scatter plot of normalized reads in HOM peaks (x axis) and SYN peaks (y axis) delineates SYN-enriched peaks (orange), SYN-depleted peaks (blue), and peaks expressed at equal levels in HOM and SYN (grey). Genes carrying the most abundant and enriched peaks at synapses are labeled. b. Peak quantification was validated by qPCR of cDNA libraries. Scatter plot of log2 fold change (PCR, x axis) and log2 fold change (RPKM, y axis) reveals high concordance. c. IGV views of selected abundant and synaptically enriched peaks. Peaks are represented as subtracted read densities (IP minus Input) in SYN (orange) and HOM (blue). d. Top 20 clusters with their representative enriched terms in SME.

High concordance between relative expression and (de)enrichment of methylation between m^6^A-seq and qPCR measurement was detected (R^2^=0.80795 for HOM and 0.87021 for SYN, Fig. 3b, Supplementary Fig. S7), validating the results of bioinformatic analysis. Inspection of individual peaks showed more prominent m^6^A peaks in SYN (Fig. 3c). Gene ontology (GO) analysis identified the most significant ontology cluster associated with SME as “central nervous system development” (Fig. 3d), demonstrating the strong functional specificity of the localized m^6^A epitranscriptome. A subset of functional clusters was converted into a network layout to illustrate their functional relationships (Supplementary Fig. S8).

Known functions of top candidates based on peak abundance in SYN were examined using manual annotation and data mining in Pubmed. Genes with the top differentially methylated peaks were associated to regulatory mechanisms of synapse formation, function, and elimination in response to local environment, mediated by neuron, astrocytes, and microglia. A recurring theme was neurological disorders such as schizophrenia, autism, anxiety, and intellectual disability (Table 1).

**Table 1:**
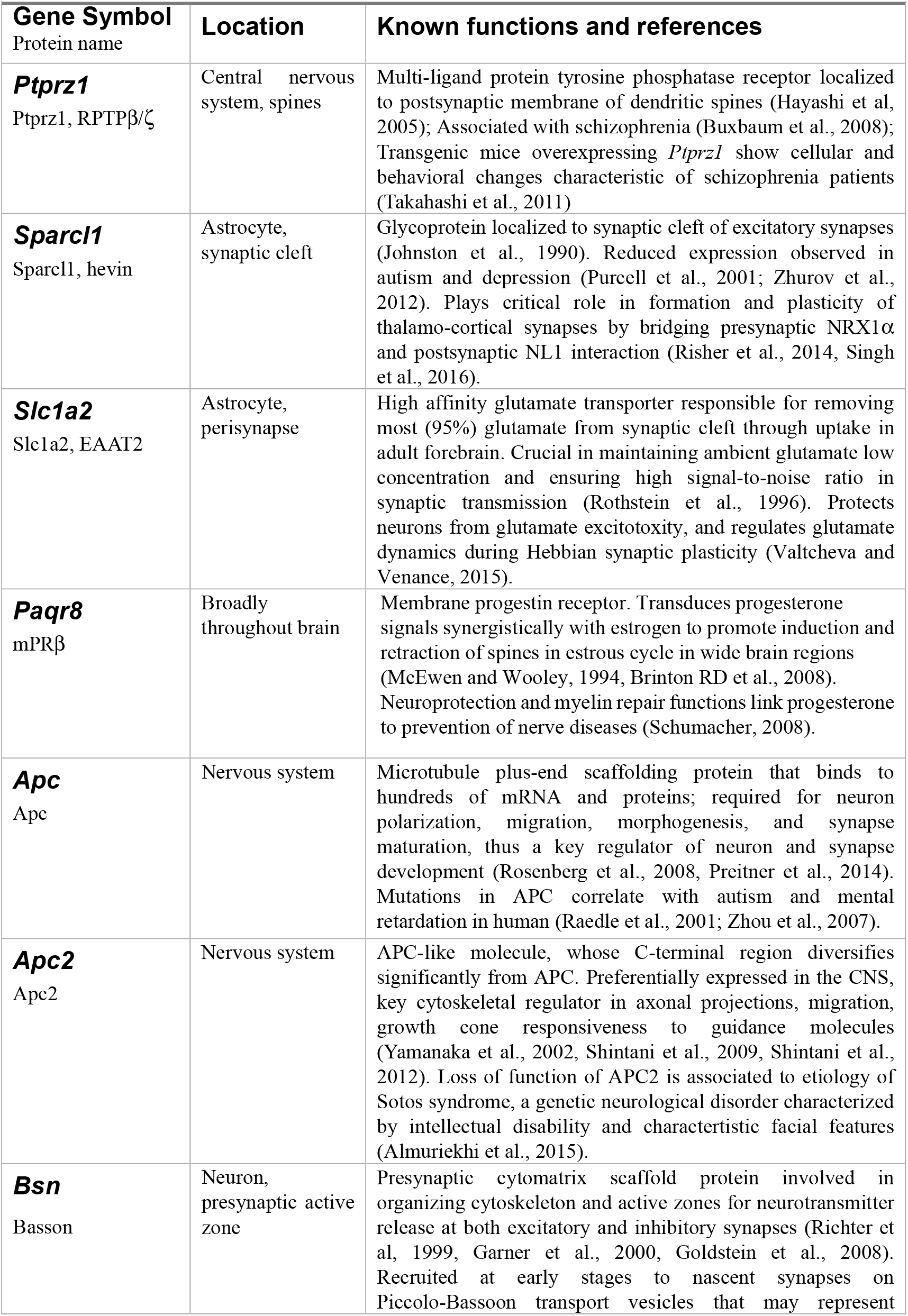
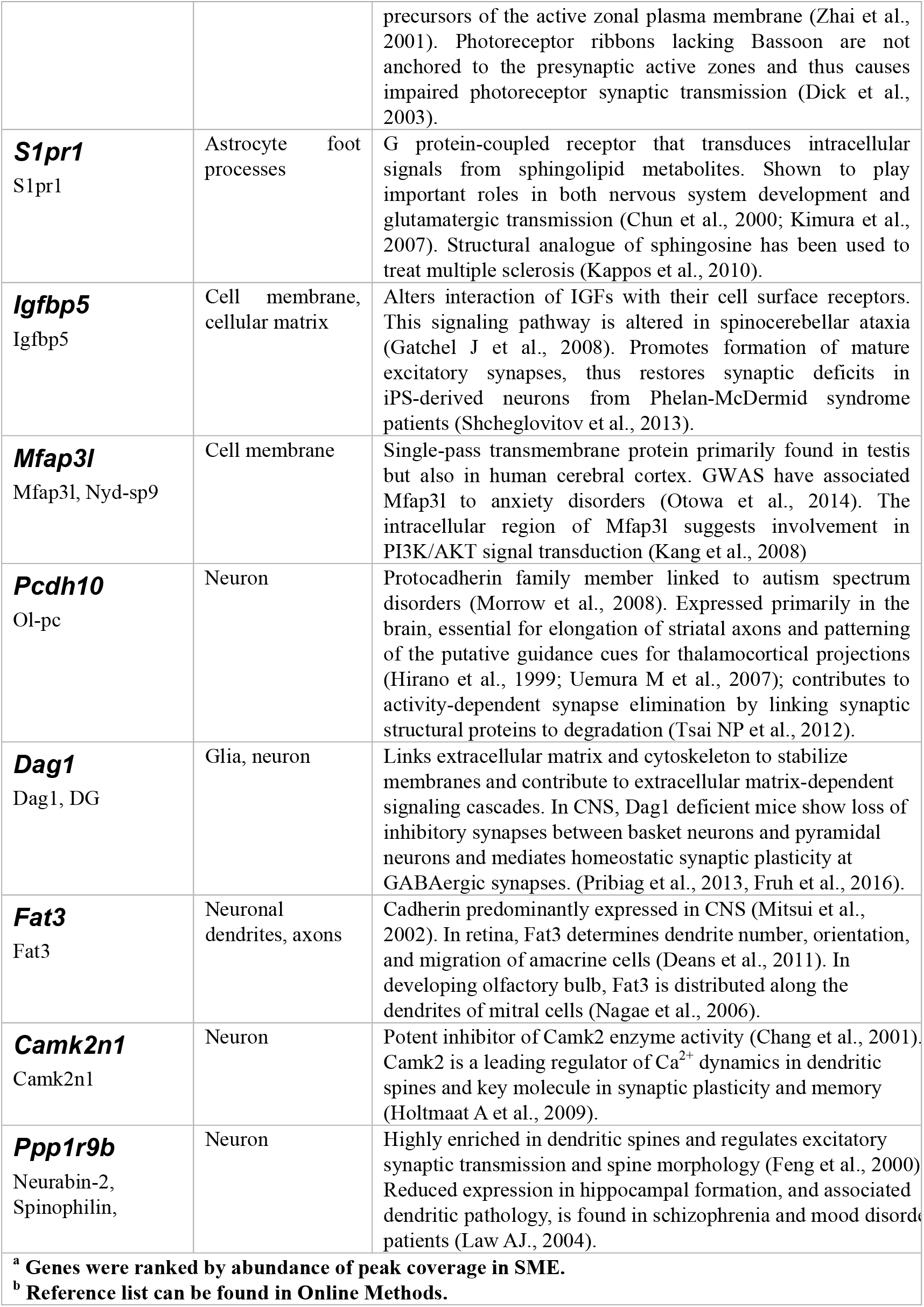
Top methylated transcripts in synaptic m^6^A epitranscriptome^30^

### Functional partitioning of synaptic transcriptome by m^6^A modification

To investigate the functional distinction between hypermethylated and hypomethylated transcripts at synapses, we compared SME to the synaptic transcriptome (ST, Fig. 1d). 50% of SME genes overlapped with ST (1,505 overlapping genes), while 1,562 non-overlapping ST-specific genes represent a hypomethylated population, and 1,482 SME-specific genes represent a synaptically hypermethylated population of transcripts (Fig. 4a and Supplementary Table S11). GO analysis revealed that the synaptically hypomethylated genes are functionally polarized to cellular metabolic processes such as “formation of a pool of free 40S subunits”, “oxidation-reduction process”, “metabolism of lipid and lipoproteins”, and “lysosome”. Transcripts enriched both by expression and methylation had greater functional applicability to CNS, such as “regulation of cell migration”, “cell morphogenesis”, and “response to growth factor”. The synaptically hypermethylated genes, however, had a striking functional delineation toward synapses, exemplified by the terms “central nervous system development”, “synapse organization”, and “anterograde trans-synaptic signaling”, all within the top 10 functional clusters with significant q-values (Fig. 4b). Furthermore, significance heat-map showed a compelling distinction between hypo- and hyper-methylated genes associated with the terms in cellular metabolism and synaptic function (Fig. 4c). These results suggest that m^6^A methylation predicts functional partitioning of transcripts at the synapse, and could thereby potentially affect the synthesis of critical synapse-specific components that alter synaptic organization and transmission (GO analyses using expressed genes as background in Supplementary Tables 12 and 13).

**Figure 4.**
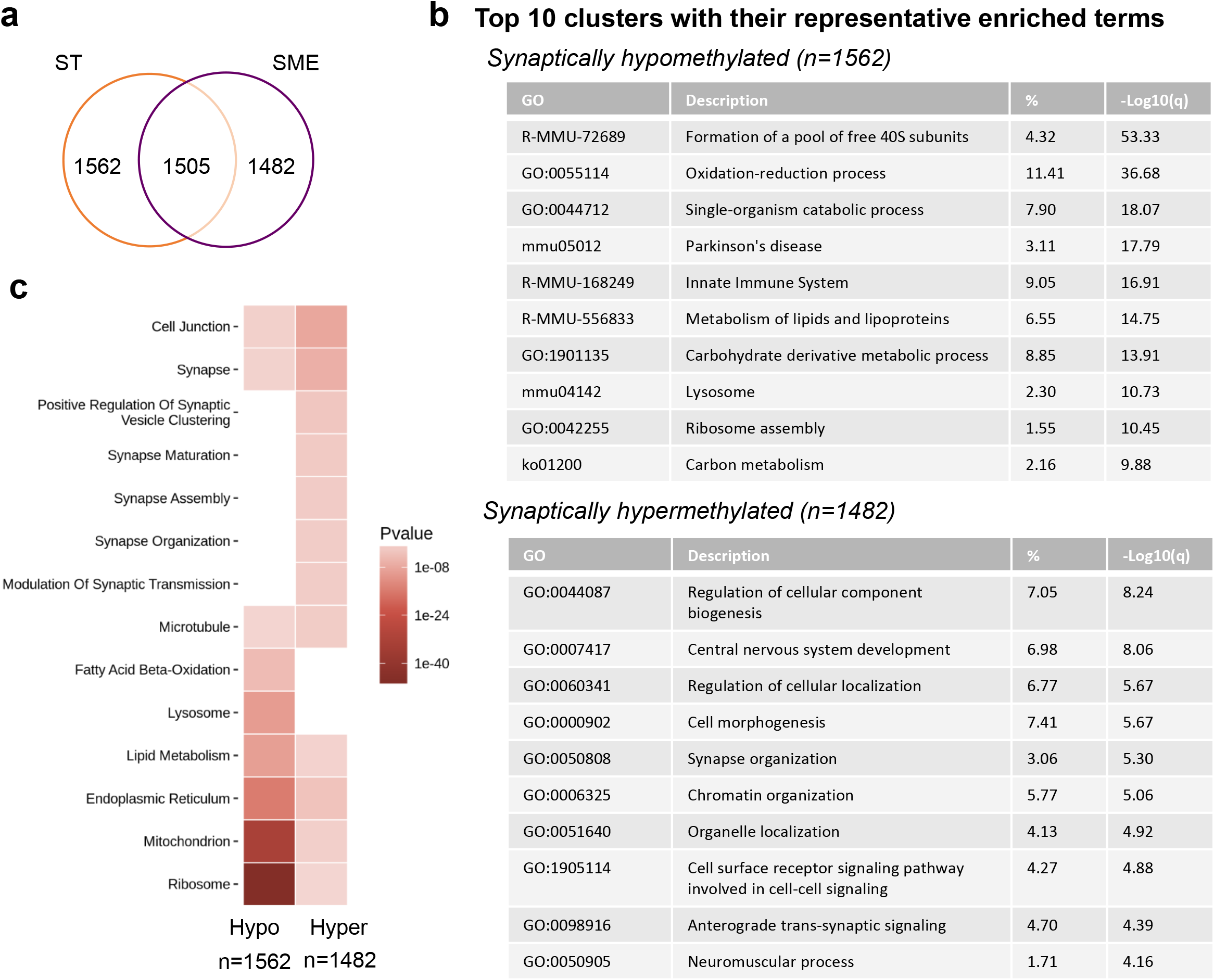
m^6^A functionally partitions synaptic transcripts for dynamic synapse formation and modulation. a. Venn diagram relating synaptic transcriptome (ST) and synaptic m^6^A epitranscriptome (SME). b. Top 10 clusters with their representative enriched terms in synaptically hypomethylated and hypermethylated gene lists. c. Significance heat-map (by p value, FDR) showing enrichment of each term category in synaptically hypo- and hypermethylated gene lists, significance cutoff, 0.05.

### Dendritic localization of m^6^A regulatory proteins

Selective enrichment of methylated transcripts at synapses raises the possibility of synapse autonomous regulatory mechanisms. We next asked whether m^6^A regulators could be detected near synapses. Using immunocytochemistry, we found wide distribution of m^6^A writers, erasers and readers in adult mouse brain slices in the cortex, thalamus, hippocampus and other brain regions. In cortical pyramidal neurons, METTL14, FTO, and YTHDF1, YTHDF2, and YTHDF3 were readily detectable in extra-somatic regions (Fig. 5a). These *in vivo* localization patterns were replicated in dissociated hippocampal neuronal cultures (Fig. 5b). Co-staining with phalloidin detected METTL14, FTO, and all three YTHDF readers in dendritic shafts adjacent to F-actin-rich spines (Supplementary Fig. S9). Particularly, all three m^6^A readers YTHDF1, YTHDF2, and YTHDF3 were localized to dendrites in a non-diffuse pattern that partially overlapped with MAP2-positive microtubules (Fig. 5c). Co-staining with presynaptic markers (synapsin I or synaptophysin) demonstrated that all three readers were localized to the vicinity of synapses. Among the three readers, poly(A) RNA colocalized the best with YTHDF1, then YTHDF3, and the least with YTHDF2 in dendrites (Fig. 5c). These results suggest that YTHDF1 may bind to m^6^A-mRNA in dendrites near synapses, thus may regulate their metabolism locally. In perfused brain slices, YTHDF1 was detected in the MAP2 positive layers, albeit the densely packed cortical environment did not resolve clear colocalization (Supplementary Fig. S10).

**Figure 5.**
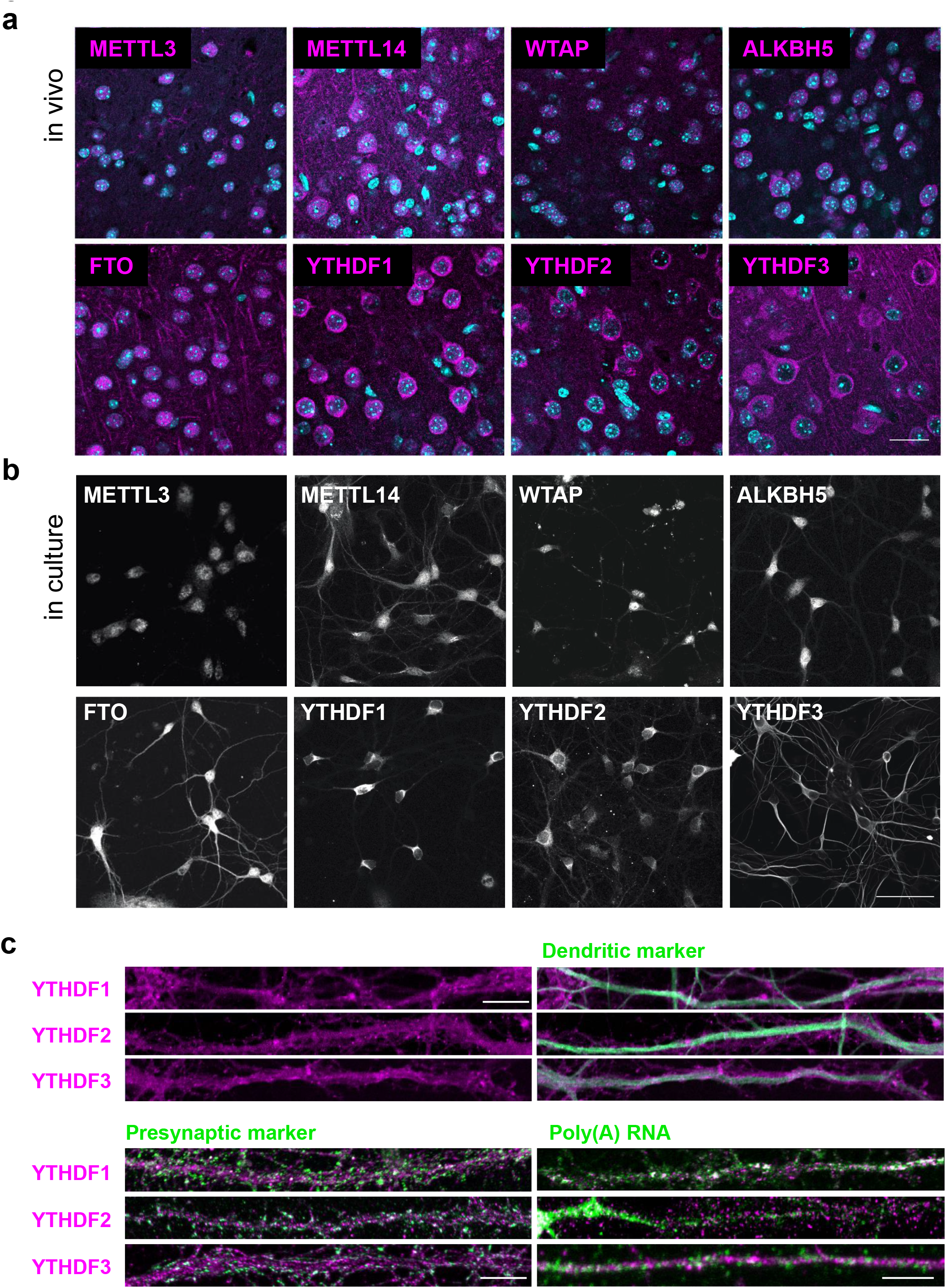
m^6^A regulatory proteins are distributed to dendrites and near synapses both in brain slices and in cultures. a. Confocal images of m^6^A writers, erasers, and readers in pyramidal cell layers of perfusion-fixed mouse cortical slices. DAPI: cyan; Immunoreactivity: magenta; Scale bar, 20μm. b. Replicate of a in cultured hippocampal neurons. Scale bar, 40μm. c. Confocal images of dendritic processes of dissociated hippocampal neuronal cultures immunostained with m^6^A readers YTHDF1–3, co-stained with dendritic marker MAP2, presynaptic marker synapsin I or synaptophysin, and poly(A) RNA. Scale bar, 3 μm.

### Reduced expression of m^6^A reader YTHDF1 causes synaptic dysfunction

We first attempted to reduce m^6^A activity by knocking down methyltransferase METTL3 in cultured neurons and observed massive cell death, which prohibited further functional analysis (Supplementary Fig. S11). We next sought to manipulate m^6^A-mediated regulatory pathways by expressing YTHDF1 shRNAs in dissociated hippocampal neurons (Fig. 6a, Supplementary Fig. S12). In YTHDF1-KD neurons, altered spine morphology, characterized by increased spine neck length (1.69±0.17 in shYTHDF1–1, 1.10±0.09μm in shYTHDF1–2, 0.73±0.06μm in shScramble) and decreased spine head width (0.32±0.01 in shYTHDF1–1, 0.31 ±0.02 μm in shYTHDF-21, 0.51 ±0.02 μm in shScramble) was detected, as well as significantly reduced PSD-95 cluster intensity (1.9-to 2.2-fold decrease in shYTHDF1–1 and shYTHDF1–2, respectively, Fig. 6b-d). Although spine density was not significantly affected, the percentage of spines containing visible PSD-95 clusters decreased (36.8% and 58.6% in shYTHDF1–1 and shYTHDF1–2, 95.0% in shScramble). In parallel experiments, expressing CRISPR/Cas9 targeting YTHDF1 in hippocampal neurons recapitulated the abnormal spine morphology (Supplementary Fig. S13), supporting specific association of the phenotypes with reduced YTHDF1 expression. These data suggest that m^6^A recognition/binding protein YTHDF1 is required for normal spine development and post-synaptic density formation.

**Figure 6.**
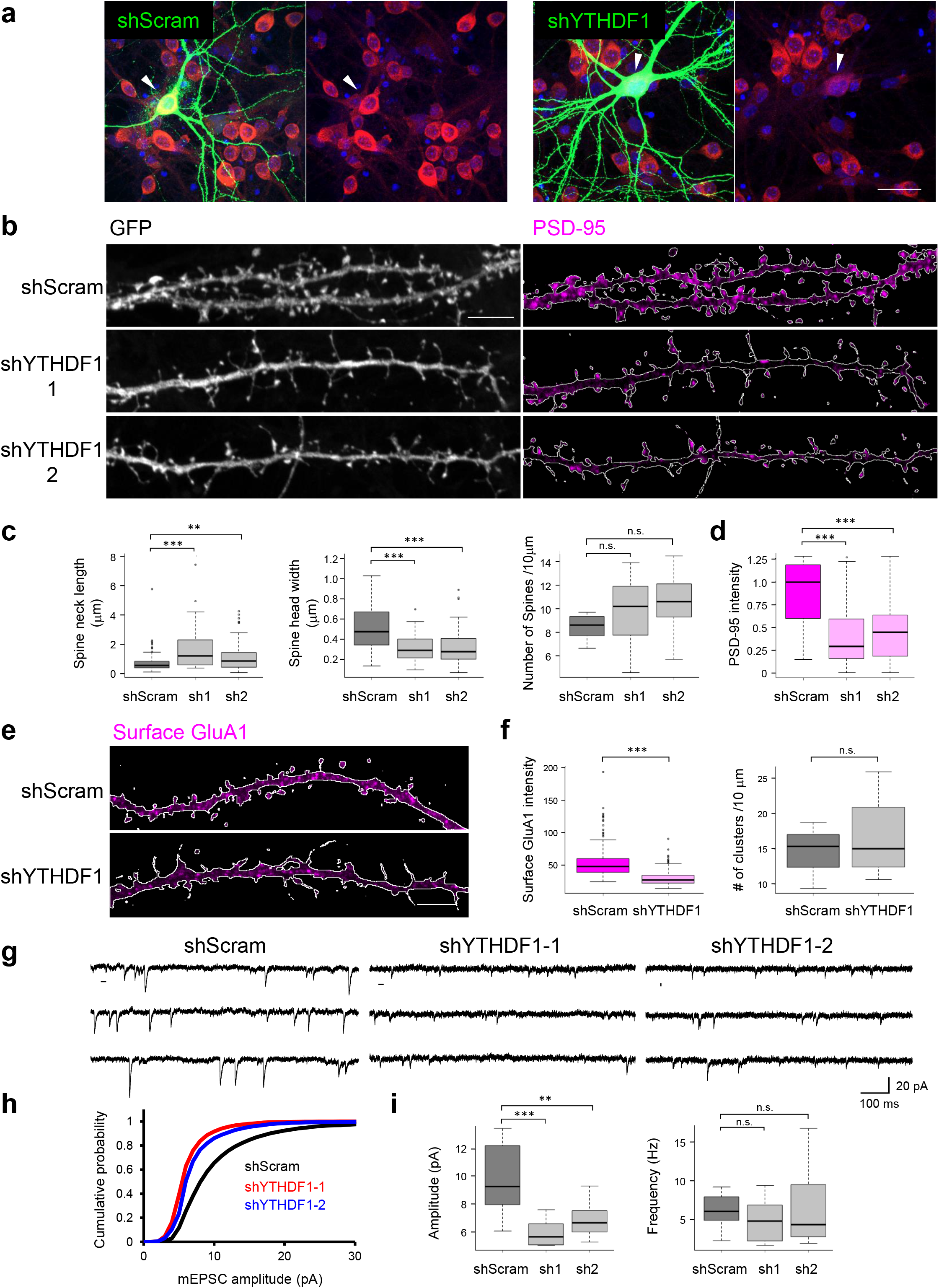
Reducing YTHDF1 expression in hippocampal neurons causes structural and functional synaptic deficits. a. Confocal images of DIV19 dissociated hippocampal neurons expressing shScramble-GFP (left panel) or shYTHDF1-GFP (right panel). DAPI (blue), GFP (green), YTHDF1 (red). Arrowheads point to GFP(+) shScram or shYTHDF1 transfected neurons. Scale bar, 10 μm. b. Confocal images of dendritic processes of DIV19 dissociated hippocampal neurons expressing shScramble-GFP or shYTHDF1-GFP (1 or 2). Left, GFP labels morphology of dendritic shaft and spines; Right, PSD-95 staining in the same samples to label post-synaptic density. Contour of affected dendrites was traced as the white lines through GFP expression. Fluorescence PSD-95 signals outside of the contour were masked to restrict quantification to the affected neurons. Scale bar, 5 μm. c. Group quantification of spine neck length, spine head diameter, and spine density; d. Group quantification of PSD-95 cluster staining intensity in spines. e. Confocal images of surface GluA1 staining in DIV20 dissociated hippocampal neurons expressing shScramble-GFP or shYTHDF1-GFP. Contour of affected dendrites is indicated by white line. Scale bar, 5 μm. f. Group quantification of fluorescence intensity of surface GluA1 cluster and GluA1 cluster density along 10μm dendrites. g. Representative mEPSC traces recorded from neurons transfected with shScramble or shYTHDF1 (sh1 or sh2). h. Cumulative plots of mEPSC amplitude distribution. i. Group quantification of amplitudes and frequencies of mEPSCs (n = 12 cells per condition).

Lack of PSD-95 clustering in spines indicated dysregulation of postsynaptic signaling complex assembly and synaptic transmission at the excitatory synapses. We therefore measured surface expression of AMPA-type glutamate receptors, which mediate fast excitatory transmission. Using an antibody recognizing extracellular N-terminus of GluA1 subunit under membrane non-permeable staining conditions, we detected significantly reduced surface expression of GluA1 (fluorescence intensity 53.4±23.7 in shScram vs 29.8± 11.0 in shYTHDF1) but unaltered cluster density (14.5±4.73 in shScram vs 16.7±6.55 clusters per 10 μm in shYTHDF1), indicating impaired trafficking of GluA1 to the synaptic membrane and/or anchoring (Fig. 6e and 6f). Such deficits can result in compromised synaptic transmission. We therefore proceeded to record miniature excitatory post-synaptic currents (mEPSCs) and found a significant reduction of amplitude (9.88±0.70 pA in shScram vs 5.92±0.27 pA and 6.86±0.35 pA in shYTHDF1–1 and shYTHDF1–2, respectively) but not of frequency (6.19±0.57 Hz in shScram vs 4.93±0.78 Hz and 6.25±1.34 Hz in shYTHDF1–1 and shYTHDF1–2, respectively, Fig. 6g-i). These results collectively demonstrate that reduced expression level of m^6^A reader YTHDF1 in neurons causes both structural and functional deficits of excitatory synapses.

### SME member APC expression is altered in YTHDF1-KD neurons

The functional deficits of synapses may be a product of altered SME expression in YTHDF1-KD neurons. YTHDF1 has been previously shown to facilitate translation of m^6^A mRNAs^43^. Toward this end, we surveyed potential candidates by crossing SME to a previously published YTHDF1 CLIP/IP dataset^43^. Among 345 common mRNA, an interesting target was APC (adenomatous polyposis coli), whose mRNA was stabilized but translational efficiency was decreased in siYTHDF1 cells^43^. Given the critical role of APC in regulating microtubule dynamics, we suspected that its altered expression in YTHDF1-KD neurons may have contributed to the synapse malfunction.

To test this possibility, we performed single-molecule staining of *Apc* mRNA simultaneously with YTHDF1 protein in cultured hippocampal neurons. GFP expression was used for single neuron morphology labeling. *Apc* mRNA was detected in dendrites including distal sites, and partially colocalized with YTHDF1 protein (Fig. 7a). In YTHDF1-KD neurons, a remarkable cell-wide decrease in APC immunoreactivity was detected, including depletion from dendrites near synapses; in contrast, APC2, another top SME candidate and APC homologue, showed unaltered expression (APC: 168.9±31.5 vs 25.4±11.3 A.U., APC2: 129.5±16.4 vs 130.6±14.5 A.U. Fig. 7b and 7c, Supplementary Fig. S14). Importantly, *Apc* mRNA level was comparable between YTHDF1-KD and control cells (10.3±9.90 vs 7.00±7.15 fluorescence puncta per cell, Fig. 7d and 7e). These data suggest that expression of localized SME members is modulated by YTHDF1 expression, exemplified by *Apc* mRNA, whose translation, but not stability or dendritic targeting, is dependent on m^6^A reader YTHDF1.

**Figure 7.**
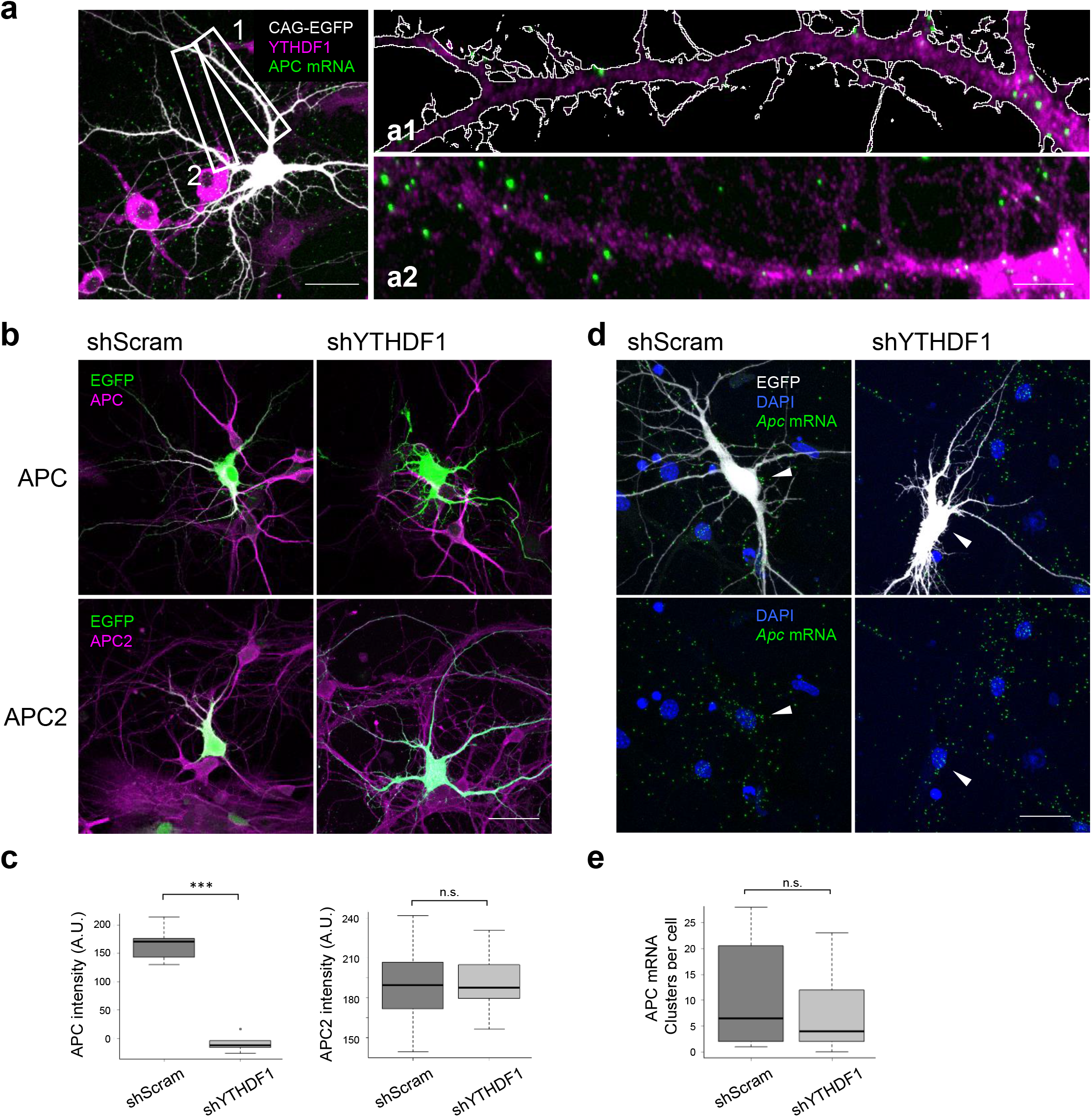
Dendritically localized SME member *Apc*, but not *Apc2* mRNA is dependent on YTHDF1 for translation. a. Confocal images of DIV7 dissociated hippocampal neurons transfected with pCAG-EGFP. Left, merged images of GFP (white), YTHDF1 immunostaining (magenta) and *Apc* mRNA (green); Right, dual-color images of YTHDF1 (magenta) and *Apc* mRNA (green) in insets a1 and a2; Contour of GFP(+) dendrites is indicated by white lines. Scale bar, 10 μm and 5 μm. b. Confocal images of DIV6 neurons with merged GFP (green) and APC (top, magenta) or APC2 (bottom, magenta). Scale bar, 10 μm. c. Group quantification of APC and APC2 immunostaining intensity in GFP(+) neurons. d. Confocal images of DIV7 dissociated hippocampal neurons transfected with shScramble-GFP or shYTHDF1-GFP (sh1) neurons with merged DAPI (blue), GFP (white) and *Apc* mRNA FISH (green). Scale bar, 10 μm. e. Quantification of *Apc* mRNA staining clusters per cell in GFP(+) neurons.

### SME highlights cell surface-associated intercellular signaling in neuroglia network

Within the SME-enriched functional terms “synapse” and “cell junction,” surface receptor pathways involved in cell-cell signaling were among the most abundant methylated transcripts. To further dissect SME function, we crossed our data set to a recently identified collection of synaptic cleft residents^44^, and found that 85% of synaptic cleft proteins at both excitatory and inhibitory synapses were encoded by methylated transcripts. Thus methylation-mediated regulation may have a wide impact on the biosynthesis of this functional class of proteins (Fig. 8a). In addition to highly abundant astrocytic methylated transcripts *Sparcl1* and *EAAT2*, whose proteins promote synapse formation and remove glutamate from synapses, we identified complement cascades in microglia surveillance in SME. The mRNAs encoding CX_3_CL1, secreted by neurons potentially as a “find me” signal, and CX_3_CR1, the receptor for CX_3_CL1 expressed exclusively by resident microglia in enhancing synapse refinement, were both highly methylated (Fig. 8b, Supplementary Fig. S15). By further data mining and assigning the annotated SME genes to different cell-types, a preliminary yet exciting tripartite SME network emerged (Fig. 8c). Critical components in all synaptic domains such as pre-synaptic protein bassoon, post-synaptic CamK2n1, a large portfolio of pre- and post-synaptic adhesion proteins, secreted proteins and reciprocal receptors involved in neuron-astrocyte and neuron-microglia communication were encoded by methylated mRNAs at the synapse, of which many have been identified as playing critical roles in sculpting synaptic connections.

**Figure 8.**
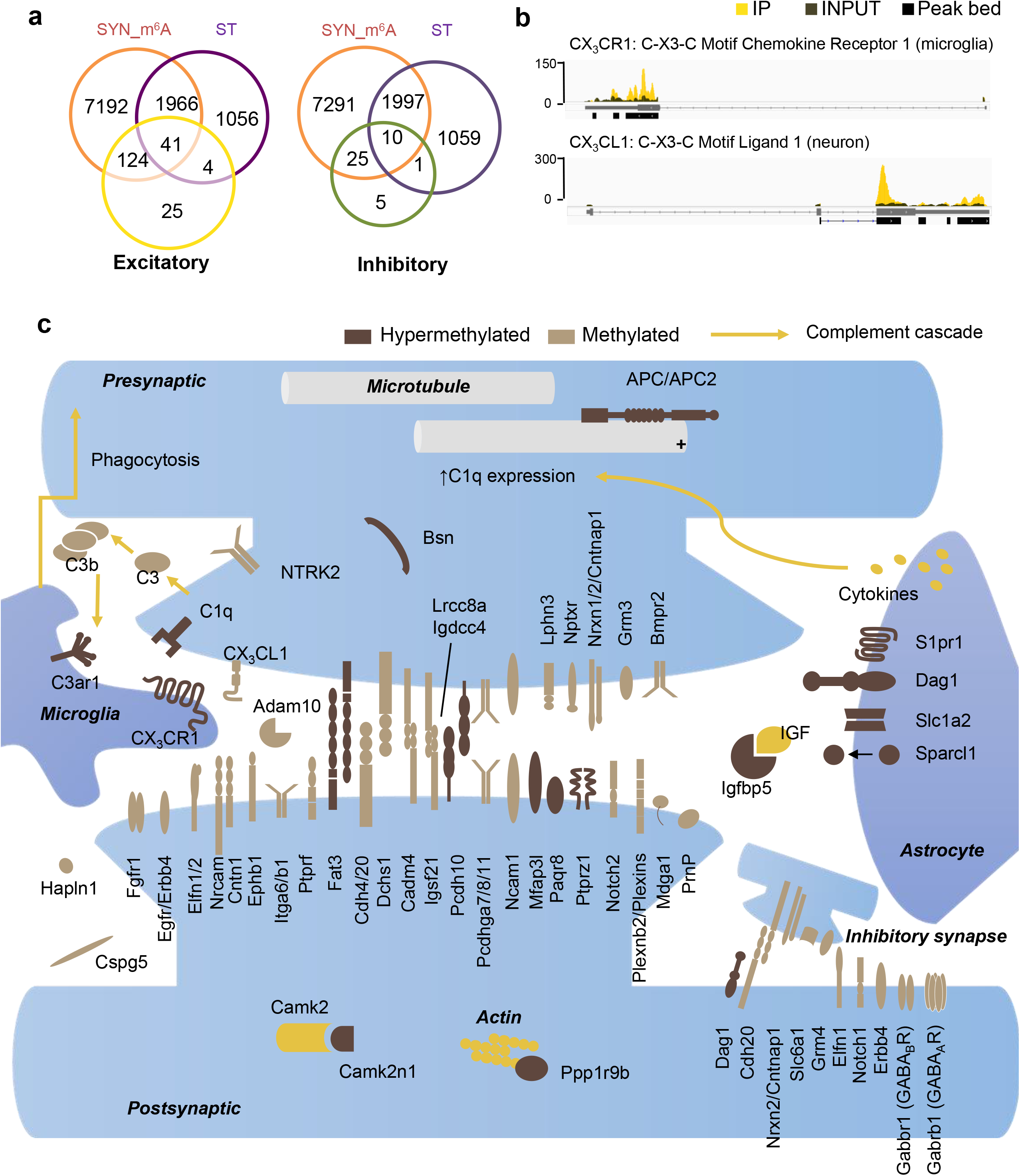
SME encapsulates surface-associated intercellular signaling network involving tripartite synapse. A. Venn diagram of methylated transcripts in SYN (SYN_m^6^A), transcripts enriched in SYN (ST), and excitatory synaptic cleft proteins (left) and inhibitory synaptic cleft proteins (right) recently identified by Loh et al., 2016. 165 of 194 (85%) excitatory, and 35 of 41 (85%) inhibitory, synaptic cleft proteins are encoded by methylated mRNA identified in SYN. B. IGV view of peak coverage of mRNAs encoding chemokine-mediated neuron-microglia communication pathway CX_3_CL1-CX_3_CR1. C. Curated assignment of proteins encoded by methylated mRNA encoding synaptic cleft proteins and by mRNAs carrying the most abundant methylation sites in SME at excitatory and inhibitory tripartite synapse.

## Discussion

The protein composition of synapses is highly dynamic and regulated in an activity-dependent manner. mRNA trafficking and local protein synthesis at a synapse plays an essential role, supplying the initial changes in protein abundance required for maintaining and modifying connections^18,45^. A rich cellular toolbox of post-transcriptional regulatory mechanisms such as microRNA, nonsense mediated decay, RNA binding proteins, and post-translational modification of translation elongation factors has been shown to regulate activity-induced translation at synapses. The SME identified in the current study reveals another layer of complexity in the regulation of the synaptic proteome. This enzyme-mediated, reversible modification, and its impact on the metabolism and translation of individual mRNA, comprises a highly flexible regulatory mechanism, effective at the single copy level.

SME consists of 4,329 m^6^A peaks that are found selectively enriched at the synapse; half of the 2,987 genes carrying these peaks are not preferentially localized to synapse. This suggests the existence of mechanisms for differential methylation or, alternatively, selective mRNA trafficking which have yet to be identified. Functional enrichment analysis revealed that the 1,482 hypermethylated genes are enriched for synapse-related functions such as synapse maturation, organization, assembly, and modulation of synaptic transmission, in sharp contrast to the localized hypomethylated transcripts, which are primarily involved in cellular metabolic pathways. The functional impetus for this deliberate partitioning is not clear. An intriguing possibility is that m^6^A-mediated spatiotemporal control of gene expression is required for coordinating activity-dependent translation of a cohort of mRNA. In support of this, a rapid and site-specific surge in m^6^A level was shown in response to UV-triggered DNA damage, indicating that changes in modification status can occur in a subset of RNAs involved in DNA repair pathways^46^. Moreover, m^6^A can exert control over translation via direct recruitment of ribosomes^47,48^ and/or co-translational interaction dynamics between the protein products^49^. Together these observations support a localized, activity-dependent dynamic proteomic network regulated by m^6^A at the synapse.

We found writer METTL14 and eraser FTO distributed to the distal processes of pyramidal neurons and, although the methylation activity of standalone METTL14 is under debate^31^, their presence raises the possibility of local enzyme-mediated (de)methylation. In future studies, this possibility could be tested using in vitro measurement of RNA methylation after stimulating isolated synapses. We also detected all three cytoplasmic readers (YTHDFs) distributed distally, suggesting versatile downstream signaling potential. Previous studies have demonstrated that binding of YTHDF1–3 to m^6^A mRNAs enhances translation efficiency and RNA degradation^43,47,48,50,51^, and in general, m^6^A is associated with shorter RNA half-life^51,52^. These apparently conflicting roles of m^6^A has led to a working model of sequential binding of YTHDF1 and YTHDF2 to help methylated mRNA achieve steady-state levels rapidly, with YTHDF3 acting as an enhancer of both degradation and translation^53,54^, thereby decreasing response time between stimulus and phenotypic output^43^. This model may also help reduce expression “noise” by degrading excess mRNA and enhancing translation of scarce transcripts. Both mechanisms can benefit synaptic stability and plasticity in response to activity, and we predict the localized readers act synergistically with the local m^6^A epitranscriptome to achieve robust production of synaptic responses.

Knockdown of reader YTHDF1 in cultured primary neurons resulted in synaptic transmission deficits and immature morphology of spines. Such phenotypes can be associated with synaptic pathophysiology. We identified a downstream molecular target of YTHDF1, SME member *Apc mRNA*, whose abundance and dendritic localization were not affected, but expression decreased with YTHDF1 knockdown. Previously, the APC cKO mouse model has shown abnormality of central synapses (increased spine density, frequency of miniature excitatory synaptic currents, and long-term potentiation), leading to a multi-syndromic neurodevelopmental disorder^55^, suggesting that altered APC expression and subsequent cytoskeletal regulation might partially contribute to the synaptic dysfunction. Dysregulated m^6^A mRNA translation at synapses may underlie disrupted long-term synaptic plasticity, learning and memory, and neurodevelopmental disorders. In support of this, human genetic research has identified the m^6^A reader YTHDC2 as one of the risk factors for autism spectrum disorder^56^. The functional analysis of synapses in the current study therefore warrants further investigation in m^6^A-mediated local translation and its relation to neuropsychiatric disorders.

A fascinating aspect of SME is its invoking of the tripartite local translation model, implicating mRNA methylation-mediated local translation in multicellular interactions between neurons, astrocytes, microglia, and oligodendrocytes. Methylated mRNA encoded proteins that coordinate intercellular cross-talk, such as postsynaptic membrane receptors (Ptprz1), synaptogenic proteins secreted from peri-synaptic astrocyte processes (Sparcl1), neuronal regulators of synapse turnover (Paqr8 and Igfbp5), chemokines secreted by neurons to attract microglia (CX_3_CL1), and the reciprocal receptors on the microglial membrane (CX_3_CR1). Advanced understanding of molecular communication pathways is required to elaborate how local translation supports combined contributions of synaptic components to the activity and plasticity of tripartite synapses. SME provides an initial candidate list for future mechanistic dissections, highly enriched in specified synaptic functions. Interestingly, of 224 recently identified peri-synaptic astrocyte mRNAs, 180 (80%) overlapped with ST and 152 (68%) overlapped with SME, in contrast to only 23% overlap between the 116 soma-derived astrocyte mRNA and ST^22^. Thus, RNA metabolism in peri-synaptic astrocyte processes may be directly regulated by m^6^A. Future molecular dissection of m^6^A regulation in tripartite synaptic function requires site-directed manipulation of the methylated adenosine residue and/or adjacent residues to disrupt methylation dynamics on individual transcripts. A recent single nucleotide-resolution m^6^A-seq method may provide valuable information in this regard^57^.

Ultimately, m^6^A may be one of many covalent RNA modifications regulating synaptic epitranscriptome^30,58^. 5mC methylase NSun2 has been shown to colocalize with FMRP in dendrites, where it may exert control over synthesis of a subset of mRNA in an activity-dependent manner^59^. NSun2 mutations have been associated with intellectual disability as well as other neurodevelopmental deficits^60,61^. The diverse functional impact of modified transcripts suggests that RNA modification can greatly enhance customization of the local proteome. Such mechanisms are perhaps indispensable for generating context-dependent responses at synapses with spatiotemporal coordination; and aberrant modification events may therefore lead to the pathogenesis of neurodevelopmental and neuropsychiatric disorders.

## METHODS

Methods and any associated references are available in the online version of the paper. Accession code PRJNA388019.

## Acknowledgements

We thank Yasunori Hayashi for critical reading of this manuscript. CeMI imaging center at iCeMS and supporting facility at medical school of Kyoto University for technical support. This work was supported by grants KAKENHI17H03546, KAKENHI26702038, KAKENHI26115515, Hirose foundation, Astellas foundation to DOW. BJG is supported by Japan Society for the Promotion of Science foreign researcher fellowship.

## Author Contributions

Wang DO conceived and designed the project. Hong WT purified synaptosomes, performed RNA extractions and m^6^A dot-blot, data mining. Wang DO and Ohara T performed library construction; Merkurjev D, Iida K, Goldie BJ, Yamaguti H, Pellegrini M performed bioinformatics analysis; Ohara T performed immunostaining; Oomoto I constructed shRNA and performed KD in cultured hippocampal neurons. Kawaguchi S performed electrophysiology. Wang DO and Goldie BJ wrote the manuscript, Martin K and Hirano T supervised parts of the project. All authors participated in data analysis and interpretation, and made indispensible contributions.

Supplementary tables 1. Results summary of cross-database comparison of m^6^A mRNA and localized mRNA lists.

Supplementary Table S2. Sequencing summary.

Supplementary Table S3. Percentage of reads removed after RNA sorting.

Supplementary Table S4. QC reads mapped with STAR to mm9 refseq.GTF

Supplementary Table S5. Differentially expressed gene lists between HOM and SYN.

Supplementary Table S6. HOM peaks and SYN peaks.

Supplementary Table S7. Methylated gene lists with peaks associated with 5’UTR, START codon, STOP codon, and 3’UTR.

Supplementary Table S8. GO analysis of genes associated with STOP codon peaks using all genes expressed either in HOM or SYN as background (supplementary to Fig. 2i).

Supplementary Table S9. GO analysis of genes associated with STOP codon peaks using genes expressed in SYN as background (supplementary to Fig 2i).

Supplementary Table S10. Synaptic m6A epitranscriptome (SME). Supplementary Table S11. Gene lists associated with synaptic transcriptome (ST, Fig. 1d) and SME.

Supplemtary Table S12. GO analysis of hypermethylated genes using SYN input genes as background (supplementary to Fig. 4).

Supplemtary Table S13. GO analysis of hypomethylated genes using SYN input genes as background (supplementary to Fig. 4).

Supplemtary Table S14. Primers used in q-PCR validation experiments.

**Supplementary Figure S1. Preparation and characterization of purified synaptosomes**. a. Synaptosome purification from adult mouse forebrains using percoll-sucrose gradients. F3 and F4 fractions were pooled as synaptosomal fraction (SYN). Homogenate lysate without fractionation was used as control for comparison (HOM); b. Relative (de)enrichment of a panel of synaptic and somatic markers probed using western blot demonstrate synapse-enrichment. c. Separation of RNA populations was confirmed by qRT-PCR examination of nuclear lncRNA Malat1, somatic RNA beta-3 tubulin and known synaptically localized transcript CaMKIIα. d. Bioanalyzer analysis of RNA integrity in prepared fractions. HOM, homogenate; Cyto, cytoplasm, F3 and F4 (SYN), synaptosomes. e. A biological replicate of m^6^A dot-blot presented in Fig. 1b.

**Supplementary Figure S2. Correlation between biological replicate sequencing libraries**. Pairwise comparison of biological replicate sequencing libraries demonstrates strong, linear correlation between replicates.

**Supplementary Figure S3. Validation of peak calling in low input m^6^A-seq**. a. hypergeometric tests on peak overlapping to three previously published databases (supplementary to fig. 2b); b. Frequency plot of motif per 5nt per peak (y axis) against distance from peak summit (x axis) in HOM.

**Supplementary Figure S4. GO analysis of m^6^A-mRNA**. Enriched human phenotypes among genes with START codon-associated SYN peaks (ToppGene). Supplementary to Fig. 2i.

**Supplementary Figure S5. IGV confirmation of peak calling**. Top panel, synapse-enriched peak in Ckap5 transcript; bottom panel, synapse-depleted peak in Lysmd4. Red, IP tracks; blue, INPUT tracks; black, peak location track. Supplementary to Fig. 2j.

**Supplementary Figure S6. Negative impact of methylation on synaptic mRNA stability and synaptic localization**. a. (left) Box plots depicting synaptic concentrations of genes in four groups with increasing methylation level. For each gene, methylation level (ML) was calculated as IP/INPUT reads ratios in each designated RNA region (full length, 5’UTR, CDS, and 3’UTR) and divided into four groups from the least methylated to the most methylated (a; ML < −2, b; −2 <= ML < 0, c; 0 <= ML < 2, d; 2 <= ML). S.TMP: transcripts per million at SYN. (right) Scatter plot and regression model labeled with slope, p-value testing the quality of model fitting, and Pearson’s correlation coefficient *r*; b. Analogous plots representing transcript enrichment at synapse compared to whole cell. Y-axis and X-axis represent relative expression values and relative methylation values, respectively. For box plots, genes were classified into one of the four groups as follows: e; rel. ML < −1, f; −1 <= rel. ML < 0, g; 0 <= rel. ML < 1, h; 1 <= rel. ML. Genes with TPM less than 1 were excluded from the analysis.

**Supplementary Figure S7. qPCR validation of synaptic (de)enriched methylation sites detected using m^6^A-seq**. Left, relative (de)enrichment estimated by qPCR; Right, relative (de)enrichment estimated by m^6^A-seq; log2 fold change (IP/INPUT) shows positive values for methylated sites and negative values for demethylated sites. Log2 fold change (IP/INPUT) values in SYN and HOM are presented together for comparison. Supplementary to Fig. 3a and 3b.

**Supplementary Figure S8**. A subset of enriched functional clusters in Fig. 3d are depicted in a network layout (Metascape). Each node represents a single term; node size is proportional to number of input genes in that term; nodes colored by p value. Terms with a similarity score > 0.3 are linked by an edge (edge thickness reflects similarity score).

**Supplementary Figure S9. m^6^A regulatory proteins in dendritic processes of dissociated hippocampal neuronal cultures**. Confocal images of m^6^A regulatory proteins (magenta) and counter-stained with phalloidin (green) to label F actin-rich spines. Scale bar, 5 μm.

**Supplementary Figure S10. m^6^A reader YTHDF1 in perfused mouse brain slices**. Top, cortical cortex; Bottom, hippocampus and CA1 and CA3 regions of hippocampus. Scale bars, 100 μm and 50 μm.

**Supplementary Figure S11. Knockdown METTL3 in dissociated hippocampal neuronal cultures induces cell death**. a. Immunofluorescence images of hippocampal neuronal cultures transfected with shMETTL3 vectors. DAPI (blue), GFP (green), METTL3 immunostaining (red). b. Decreased METTL3 protein expression was detected within a day after transfection by immunofluorescence staining.

**Supplementary Figure S12. Knock-down efficiency estimation of YTHDF1-sh1 and YTHDF1-sh2**. a. Normalized YTHDF1 mRNA expression in shScramble, sh1, and sh2 cells (normalized to beta actin mRNA); b. Western-blots of YTHDF1 proteins in shScramble, sh1, and sh2 cells lysates. β-actin was blotted on the same membrane as loading control. Supplementary to Fig. 6.

**Supplementary Figure S13. Replicated neuron phenotypes of YTHDF1-KD cells using CRISPR/Cas9 technology**. a. GFP fluorescence images of control cells (expressing pCAG-EGFP and pX330) and YTHDF1-KD cells (expressing pCAG-EGFP and pX330-guide RNA sequences g1 or g2). Scale bar, 5 μm; b. Group quantifications of spine neck length and spine head width showed similar phenotypes to YTHDF1-KD using shRNA vectors. Supplementary to Fig. 6.

**Supplementary Figure S14. APC protein expression was altered in YTHDF1-KD neurons**. a. Confocal immunofluorescence images of APC using APC N-terminus antibody (top) or APC c-terminus antibody (bottom). GFP labels neurons expressing the shRNAs (shScramble or shYTHDF1). Both antibodies consistently detected decreased APC protein expression in YTHDF1-KD neurons; DAPI (blue), GFP (green), APC protein (magenta). Scale bar, 10 μm.

**Supplementary Figure S15. IGV view of peaks in transcripts involved in neuron-microglia signaling at tripartite synapse**. Red, synapse tracks; blue, homogenate tracks; black, peak location track. In support of Figure 8b.

## References (Harvard Style)

1. Kelsch, W., Lin, C.-W. & Lois, C. Sequential development of synapses in dendritic domains during adult neurogenesis. Proc. Natl. Acad. Sci. 105, 16803–16808 (2008).

2. Holtmaat, A. & Svoboda, K. Experience-dependent structural synaptic plasticity in the mammalian brain. Nat. Rev. Neurosci. 10, 647–658 (2009).

3. Citri, A. & Malenka, R. C. Synaptic Plasticity: Multiple Forms, Functions, and Mechanisms. Neuropsychopharmacology 33, 18–41 (2008).

4. Kang, H. et al. A requirement for local protein synthesis in neurotrophin-induced hippocampal synaptic plasticity. Science 273, 1402–6 (1996).

5. Bagni, C., Mannucci, L., Dotti, C. G. & Amaldi, F. Chemical stimulation of synaptosomes modulates α-Ca2+/calmodulin-dependent protein kinase II mRNA association to polysomes. J Neurosci 20, RC76 (2000).

6. Miller, S. et al. Disruption of dendritic translation of CaMKIIα impairs stabilization of synaptic plasticity and memory consolidation. Neuron 36, 507–519 (2002).

7. Ju, W. et al. Activity-dependent regulation of dendritic synthesis and trafficking of AMPA receptors. Nat. Neurosci. 7, 244–53 (2004).

8. Sutton, M. A. & Schuman, E. M. Dendritic Protein Synthesis, Synaptic Plasticity, and Memory. Cell 127, 49–58 (2006).

9. Ashraf, S. I., Mcloon, A. L., Sclarsic, S. M. & Kunes, S. Synaptic protein synthesis associated with memory is regulated by the RISC pathway in Drosophila. Cell 124, 191–205 (2006).

10. Mameli, M., Balland, B., Luján, R. & Lüscher, C. Rapid synthesis and synaptic insertion of GluR2 for mGluR-LTD in the ventral tegmental area. Science 317, 530–533 (2007).

11. Schmid, A. et al. Activity-dependent site-specific changes of glutamate receptor composition in vivo. Nat. Neurosci. 11, 659–666 (2008).

12. Wang, D. O. et al. Synapse- and Stimulus-Specific Local Translation During Long-Term Neuronal Plasticity. Science 324, 1536–1540 (2009).

13. Younts, T. J. et al. Presynaptic Protein Synthesis Is Required for Long-Term Plasticity of GABA Release. Neuron 92, 479–492 (2016).

14. Bassell, G. J. & Warren, S. T. Fragile X Syndrome: Loss of Local mRNA Regulation Alters Synaptic Development and Function. Neuron 60, 201–214 (2008).

15. Kelleher, R. J. & Bear, M. F. The Autistic Neuron: Troubled Translation? Cell 135, 401–406 (2008).

16. Darnell, J. C. et al. FMRP Stalls Ribosomal Translocation on mRNAs Linked to Synaptic Function and Autism. Cell 146, 247–261 (2011).

17. Santini, E. et al. Exaggerated translation causes synaptic and behavioural aberrations associated with autism. Nature 493, 411–415 (2013).

18. Cajigas, I. J. et al. The local transcriptome in the synaptic neuropil revealed by deep sequencing and high-resolution imaging. Neuron 74, 453–66 (2012).

19. Perea, G., Navarrete, M. & Araque, A. Tripartite synapses: astrocytes process and control synaptic information. Trends Neurosci. 32, 421–31 (2009).

20. Eroglu, C. & Barres, B. A. Regulation of synaptic connectivity by glia. Nature 468, 223–231 (2010).

21. Wu, Y., Dissing-Olesen, L., MacVicar, B. A. & Stevens, B. Microglia: Dynamic Mediators of Synapse Development and Plasticity. Trends Immunol. 36, 605–613 (2015).

22. Sakers, K. et al. Astrocytes locally translate transcripts in their peripheral processes. Proc. Natl. Acad. Sci. U. S. A. 71795 (2017). doi:10.1073/pnas.1617782114

23. Kiebler, M. a & Bassell, G. J. Neuronal RNA granules: movers and makers. Neuron 51, 685–90 (2006).

24. Bramham, C. R. & Wells, D. G. Dendritic mRNA: transport, translation and function. Nat. Rev. Neurosci. 8, 776–789 (2007).

25. Martin, K. C. & Ephrussi, A. mRNA Localization: Gene Expression in the Spatial Dimension. Cell 136, 719–730 (2009).

26. Wang, D. O., Martin, K. C. & Zukin, R. S. Spatially restricting gene expression by local translation at synapses. Trends Neurosci. 33, 173–82 (2010).

27. Fernandez-moya, S. M., Bauer, K. E. & Kiebler, M. A. Meet the players: local translation at the synapse. Front. Mol. Neurosci. 7, 1–6 (2014).

28. Sephton, C. F. & Yu, G. The function of RNA-binding proteins at the synapse: Implications for neurodegeneration. Cell. Mol. Life Sci. 72, 3621–3635 (2015).

29. Gilbert, W. V, Bell, T. A. & Schaening, C. Messenger RNA modifications: Form, distribution, and function. Science 352, 1408–1412 (2016).

30. Nainar, S., Marshall, P. R., Tyler, C. R., Spitale, R. C. & Bredy, T. W. Evolving insights into RNA modifications and their functional diversity in the brain. Nat. Neurosci. 19, 1292–1298 (2016).

31. Wang, Y. & Zhao, J. C. Update: Mechanisms Underlying N6-Methyladenosine Modification of Eukaryotic mRNA. Trends Genet. 32, 763–773 (2016).

32. Liu, N. et al. N6-methyladenosine-dependent RNA structural switches regulate RNA–protein interactions. Nature 518, 560–564 (2015).

33. Meyer, K. D. et al. Comprehensive Analysis of mRNA Methylation Reveals Enrichment in 3’ UTRs and near Stop Codons. Cell 149, 1635–1646 (2012).

34. Hess, M. E. et al. The fat mass and obesity associated gene (Fto) regulates activity of the dopaminergic midbrain circuitry. Nat. Neurosci. 16, 1042–8 (2013).

35. Fustin, J.-M. et al. RNA-methylation-dependent RNA processing controls the speed of the circadian clock. Cell 155, 793–806 (2013).

36. Widagdo, X. J. et al. Experience-Dependent Accumulation of N6-Methyladenosine in the Prefrontal Cortex Is Associated with Memory Processes in Mice. J. Neurosci. 36, 6771–6777 (2016).

37. Lence, T. et al. m6A modulates neuronal functions and sex determination in Drosophila. Nature 540, 242–247 (2016).

38. Walters, B. J. et al. The Role of The RNA Demethylase FTO (Fat Mass and Obesity-Associated) and mRNA Methylation in Hippocampal Memory Formation. Neuropsychopharmacology 42, 1502–1510 (2017).

39. Dominissini, D., Moshitch-Moshkovitz, S., Salmon-Divon, M., Amariglio, N. & Rechavi, G. Transcriptome-wide mapping of N6-methyladenosine by m6A-seq based on immunocapturing and massively parallel sequencing. Nat. Protoc. 8, 176–89 (2013).

40. Ke, S. et al. A majority of m6A residues are in the last exons, allowing the potential for 3’ UTR regulation. Genes Dev. 29, 1–17 (2015).

41. Schwartz, S. et al. Perturbation of m6A Writers Reveals Two Distinct Classes of mRNA Methylation at Internal and 5’ Sites. Cell Rep. 8, 284–96 (2014).

42. Dominissini, D. et al. Topology of the human and mouse m6A RNA methylomes revealed by m6A-seq. Nature 485, 201–6 (2012).

43. Wang, X. et al. N6-methyladenosine Modulates Messenger RNA Translation Efficiency. Cell 161, 1388–1399 (2015).

44. Loh, K. H. et al. Proteomic Analysis of Unbounded Cellular Compartments: Synaptic Clefts. Cell 166, 1295–1307.e21 (2016).

45. Niere, F. et al. Analysis of Proteins That Rapidly Change Upon Mechanistic/Mammalian Target of Rapamycin Complex 1 (mTORC1) Repression Identifies Parkinson Protein 7 (PARK7) as a Novel Protein Aberrantly Expressed in Tuberous Sclerosis Complex (TSC). Mol. Cell. Proteomics 15, 426–444 (2016).

46. Xiang, Y. et al. RNA m6A methylation regulates the ultraviolet-induced DNA damage response. Nature 543, 573–576 (2017).

47. Meyer, K. D. et al. 5’ UTR m6A Promotes Cap-Independent Translation. Cell 163, 999–1010 (2015).

48. Zhou, J. et al. Dynamic m6A mRNA methylation directs translational control of heat shock response. Nature 526, 591–594 (2015).

49. Choi, J. et al. N6-methyladenosine in mRNA disrupts tRNA selection and translation-elongation dynamics. Nat. Struct. Mol. Biol. 23, 110–115 (2016).

50. Du, H. et al. YTHDF2 destabilizes m6A-containing RNA through direct recruitment of the CCR4–NOT deadenylase complex. Nat. Commun. 7, 12626 (2016).

51. Wang, X. et al. N6-methyladenosine-dependent regulation of messenger RNA stability. Nature 505, 117–20 (2014).

52. Fu, Y., Dominissini, D., Rechavi, G. & He, C. Gene expression regulation mediated through reversible m6A RNA methylation. Nat. Rev. Genet. 15, 293–306 (2014).

53. Shi, H. et al. YTHDF3 facilitates translation and decay of N6-methyladenosine-modified RNA. Cell Res. 27, 315–328 (2017).

54. Li, A. et al. Cytoplasmic m6A reader YTHDF3 promotes mRNA translation. Cell Res. 27, 444–447 (2017).

55. Mohn, J. L. et al. Adenomatous polyposis coli protein deletion leads to cognitive and autism-like disabilities. Mol. Psychiatry 19, 1133–1142 (2014).

56. Liu, X. et al. Genome-wide Association Study of Autism Spectrum Disorder in the East Asian Populations. Autism Res. 9, 340–349 (2016).

57. Linder, B. et al. Single-nucleotide-resolution mapping of m6A and m6Am throughout the transcriptome. Nat Methods 12, 767–772 (2015).

58. Lacoux, C. et al. BC1-FMRP interaction is modulated by 2’-O-methylation: RNA-binding activity of the tudor domain and translational regulation at synapses. Nucleic Acids Res. 40, 4086–4096 (2012).

59. Hussain, S. & Bashir, Z. I. The epitranscriptome in modulating spatiotemporal RNA translation in neuronal post-synaptic function. Front. Cell. Neurosci. 9, 1–10 (2015).

60. Abbasi-moheb, L. et al. Mutations in NSUN2 Cause Autosomal-Recessive Intellectual Disability. Am. J. Hum. Genet. 90, 847–855 (2012).

61. Roundtree, I. A., Evans, M. E., Pan, T. & He, C. Dynamic RNA Modifications in Gene Expression Regulation. Cell 169, 1187–1200 (2017).

